# Genome-wide association study revealed loci linked to post-drought recovery and traits related to persistence of smooth bromegrass (*Bromus inermis*)

**DOI:** 10.1101/2021.05.16.444329

**Authors:** Fatemeh Saeidnia, Mohammad Mahdi Majidi, Aghafakhr Mirlohi

## Abstract

Association analysis has been proved as a powerful tool for genetic dissection of complex traits. This study was conducted to identify marker–trait associations for recovery, persistence, and as well as finding stable associations. In this study, a diverse panel of polycross derived progenies of smooth bromegrass was phenotyped under normal and water stress, during three consecutive years. Association analysis was performed between nine important agronomic traits along with three seasonal growth activity indices based on 535 SRAP markers. Population structure analysis identified five main subpopulations possessing significant genetic differences. Association analysis using mixed linear mode1 identified 339 and 233 marker-trait associations under normal and water stress environments, respectively. Some of these markers were associated with more than one trait; which can be attributed to pleiotropic effects or to a number of tightly linked genes affecting several traits. If the effectiveness of these markers in genetic control of these traits is validated, they could be potentially used for initiation of marker-assisted selection and targeted trait introgression of smooth bromegrass under normal and water stress environments.

**Highlight:** In this study, stable marker-trait associations (MTAs) between years and moisture regimes (normal and water stress environments) were identified in a diverse panel of polycross derived progenies of smooth bromegrass.

## Introduction

Under the climatic changing context, drought is becoming most significant and acute problem affecting growth, survival and persistence of crops in many regions of the world, particularly in arid and semi-arid regions (Mollasadeghi *et al*., 2011; Hussain *et al*., 2012). Developing drought-tolerant varieties is an important objective of plant breeding programs and is expected to be a key component in strategies to mitigate climate change and minimize losses due to stresses and ensure production stability (Gustafson, 2011; Öztürk *et al*., 2014). Given the advantages that perennial forage grasses such as smooth bromegrass have in the arenas of both agricultural and forage sustainability for animal farming, they can be a valuable substitute for annuals under drought conditions. In these species, forage yield, persistence and recovery are of the most important agronomic features in areas with periodic drought in summer (Annicchiarico *et al*., 2011; Pecetti *et al*., 2011).

Successful adoption of perennial forage species depends on their ability to both survive through repeated summer droughts as well as maintain forage productivity (Annicchiarico *et al*., 2011; Volaire, 2008). In the other words, persistence of perennial grasses is dependent on the ability of the plant to maintain a viable crown at the soil surface from which growth regenerates. When genotypes are spaced planted, the difference in performance throughout the years can give an idea of persistence (Saeidnia *et al*., 2017b). However, it is also known that poor regrowth can result in low persistence (Nie *et al*., 2008). Therefore, during drought, trait selection for high recovering ability may be of more economic significance than selecting for improved growth (Hung and Wang, 2005; Chai *et al*., 2010). This will enables forage plants to persist in swards or pastures and to improve their competition with less drought-tolerant species (Kanapeckas *et al*., 2008; Volaire *et al*., 2014). The potential of a plant to recommence growth and grain yield after experiencing water stress is defined as drought recovery (Fang and Xiong, 2015). Therefore, successful post-drought recovery is contingent on a variety of mechanisms, such as compensatory growth of remaining tissues, retention of intact growing points throughout water stress, and mobilization of the organism’s carbohydrate reserves (Chai *et al*., 2010), and is the best criterion for drought tolerance along with the persistence.

Besides, plant survival and recovery have been associated with summer dormancy, dehydration tolerance in surviving tissues, extensive root system and the ability of roots to extract water at low soil water potentials (Norton *et al*., 2006; Shaimi *et al*., 2009). Summer dormancy is an endogenously and temporary suspension of visible growth of any plant structure containing a meristem (Norton *et al*., 2006), which leads to the reduction or cessation of leaf growth and possible senescence of herbage expressed under non-limiting irrigation in summer (Norton *et al*., 2008). Shaimi *et al*., (2009) showed that, in orchardgrass, summer dormancy is associated with superior persistence and recovery after severe drought under arid and semi-arid conditions.

Drought tolerance is a complex quantitative trait, involving diverse and multiple molecular and physiological mechanisms, signal transduction and metabolic pathways (Shinozaki and Yamaguchi-Shinozaki, 2007). Therefore, a promising strategy to facilitate selection and breeding for drought tolerance is to identify simply inherited genetic markers linked to the traits related to drought tolerance such as drought survival, post-drought recovery, and persistence. The basic prerequisite for marker-assisted selection (MAS) is the availability of markers that are strictly associated to genes or QTLs which can be used to dissect complex traits (Ebrahimi *et al*., 2017; Kempf *et al*., 2017).

The first step of MAS, as an important tool for accelerating the rate of genetic gain, is to dissect marker-trait associations (MTAs) (Moose and Mumm, 2008). Genome-wide association studies (GWAS) or association mapping, which is also known as linkage disequilibrium (LD) mapping, have recently been proved to be useful and powerful alternative to bi-parental QTL mapping for identifying MTA in plant populations (Thomson, 2014; Patel *et al*., 2015). Compared to linkage mapping, GWAS offers higher mapping resolution, is less time consuming, requires fewer resources, and evaluates a much larger gene pool rapidly (Zhu *et al*., 2008; Korte and Farlow, 2013). The power of association analysis to identify and characterize loci associated with complex traits is highly affected by admixtures of populations (Zhang *et al*., 2012). Therefore, in order to avoid identifying false positive or spurious associations between markers and traits, it is necessary to evaluate the population structure (Pritchard *et al*., 2000). In addition, utilizing a mixed-model approach involving multiple levels of relatedness simultaneously, has an important role in avoidance of both types of error (types I and II) (Yu *et al*., 2006; Ebrahimi *et al*., 2017). Association analysis can be performed by using general linear model (GLM) and mixed linear model (MLM). In MLM, both the kinship matrix (K) and population structure (Q) are incorporated, whereas in the GLM, only population structure information is used as a covariate (Ebrahimi *et al*., 2017).

In recent years, association analysis in forage grasses is applied extensively for dissecting the genetic bases of different quantitative traits in several species such as perennial ryegrass (*Lolium perenne*) (Auzanneau *et al*., 2011; Yu *et al*., 2011; Tang *et al*., 2013; Yu *et al*., 2013), tall fescue (*Festuca arundinacea*) (Lou *et al*., 2015; Sun *et al*., 2015), and orchardgrass (*Dactylis glomerata* L.) (Yan *et al*., 2016; Zhao *et al*., 2017; Abtahi *et al*., 2018b). These studies have demonstrated that GWAS is an efficient technique for identifying genomic regions linked to the quantitative traits. However, in smooth bromegrass the use of association analysis in identifying links between genes or markers with complex traits such as those related to persistence and recovery after drought is still in its infancy. To the best of our knowledge, this study is the first report in this regard. Therefore, this was conducted to: i) identify genetic loci associated with the productivity, persistence, post-drought recovery, and summer dormancy under normal and water stress environments; and ii) discover stable marker loci linked to the stated traits, between moisture environments and years.

## Materials and Methods

### Plant materials and field evaluations

Genetic materials used in this study consisted of 216 clones randomly selected from a large nursery comprised of 1800 single spaced-plant polycrossed progenies resulting from 25 preliminary parental ecotypes of smooth bromegrass (*Bromus inermis*). The 25 parental genotypes of the polycross population were collected from different regions (mainly Iran) and established in the field (Supplementary Table S1). Polycross seeds from the 25 parents were grown in plastic boxes in a greenhouse during the winter of 2010. Established seedlings were space planted in the field in first March 2010 and grown during 2011 and 2012. For the present study, 216 clones were randomly selected from the nursery, propagated in a greenhouse during winter 2012 and transferred to another field experiment, according to a randomized complete block design with 12 replications (Six replications for each normal and water stress environments) in March 2012. Spaced plants in each block were grown with 50-cm spacing between and within the rows.

Genotypes were evaluated under normal and water stress environments during 2013-2016, in which irrigation was occurred when 50% and 85% of the total available soil water was depleted from the root zone, respectively, following accepted methods of determination of evapotranspiration (Allen *et al*., 1998). Water stress was continuously applied in each year of the experiment from early May to early October (plants growing period). In this period, depending on the weather conditions, the irrigation intervals were variable during the growing season and between the two moisture environments. To determine the gravimetric soil– water content and detect the irrigation times, soil samples were daily taken from different sites of each moisture environment before irrigation at depths of 0–20, 20–40, and 40–60 cm, using a hand auger (Clarke Topp *et al*., 2008). The irrigation depth was calculated by the following formula:

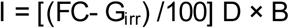

where I is the irrigation depth (cm), FC is the soil gravimetric moisture percentage at the field capacity, G_irr_ is the soil gravimetric moisture percentage at the time of irrigation, D is the root-zone depth, and B is the soil bulk density at root-zone (1.4 g cm^-3^). A basin irrigation system was used for watering. In this system, water was delivered to the field via a pump station and polyethylene pipes. A volumetric counter was used to measure water volume applied under each moisture environment.

### Phenotyping

During plant establishment year no data were recorded. Traits were measured for three years during the growing seasons of 2013 to 2015. In these years, when flowering in all plots was completed (about early summer), the aboveground biomass of each plant was harvested manually from 5 cm aboveground, dried at 75°C for 48 h and then dry matter yield weight per plant was recorded. In each year of the experiment, two harvests of above-ground biomass were undertaken. The first harvest was done after pollination assessment in late spring (spring forage yield; SPFY, DMY1), and the second in late summer (summer forage yield; SUFY, DMY2) to assess complete growth. To evaluate the seasonal growth activity, summer dormancy index (SDI) was calculated as the ratio of the SUFY of a genotype to the SPFY of the same genotype as follows (Norton *et al*., 2008):

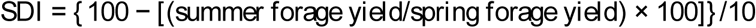

After three years of field evaluation, all genotypes were assessed for post-drought recovery in the field in 2016. For this purpose, after the first harvest, a severe water stress was imposed on both previous moisture environments (normal and water stress) by stopping irrigation for 60 days (from 1^st^ June to 31^st^ July) until complete desiccation of the grass foliage. Then to allow for water stress recovery, all plants were subsequently irrigated to the point of field capacity every week. After six weeks of regular re-watering, traits related to recovery were measured. Recovery yield (RY) was obtained by measuring the above-ground biomass of each genotype after withholding irrigation and re-watering. The degree of recovery after drought (DRAD) was visually scored based on a scale of 0 to 9. In this respect, green and fully hydrated leaves were rated as 9 and desiccated brown/dead leaves were graded as zero. Persistence (PER) of genotypes was calculated as the difference of dry matter yield of the first cut at the fourth year (2016) from dry matter yield of the first cut at second year (2014) (Saeidnia *et al*., 2017b):

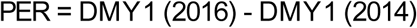

### Genotyping

Young leaf tissues of smooth bromegrass plants was used for extraction of genomic DNA, using the modified method described by Murray and Thompson (1980). The quality and quantity of DNA were determined by electrophoresis in 1% agarose gel. Genotyping using sequence related amplified polymorphism (SRAP) markers was performed following the method of Li and Quiros (2001). Among the SRAP markers available, 30 primer combinations were screened by polymerase chain reaction (PCR). PCR reactions were conducted in a final volume of 10 µL consisted of 1.5 µL of DNA, 1 µL of forward primer, 1 µL of reverse primer, 5 µL of master mix (Amplicon) and 1.5 µL of distilled water, using a BIO-RAD thermocycler. For SRAP analysis, samples were subjected to the following thermal profile: the five cycles including initial denaturation of 1 min at 94°C, annealing of 1 min at 35°C, and extension of 1 min at 72°C, followed by 35 cycles of 3 min at 50°C, with a final extension of 10 min at 72°C. PCR products were separated by electrophoresis on 12% non-denatured polyacrylamide gels and stained by AgNO3 solution (Bassam et al., 1991). Polymorphic SRAP markers were scored as binary data with presence (1) or absence (0).

### Statistical analyses

The Kolmogorov–Smirnov and Bartlett’s tests were used to examine the normality and homogeneity of variance, respectively. Analysis of variance and estimation of variance components for the normal and water stress environments separately were performed for all measured traits using PROC Mixed in SAS release 9.4 (SAS Institute, 2011). Least significant difference (LSD) test at P□0.05 was used for trait means comparison (Steel and Torrie, 1980). Broad-sense heritability (h^2^_b_) was estimated for normal and water stress environments on a phenotypic mean basis averaged over replications as described by Nguyen and Sleper (1983):

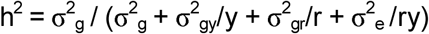

Where *σ*^*2*^ _*g*_ is the genotype, *σ*^*2*^ _*gy*_ is the genotype × year, *σ*^*2*^ _*rg*_ is the genotype × rep and *σ*^*2*^ _*e*_ is the residual variance, y is the number of years and r is the number of replicates. To estimate the level of genetic variation, the phenotypic coefficient of variation (PCV) and genetic coefficient of variation (GCV) were calculated as:

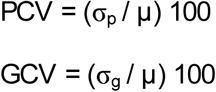

where σp is the standard deviation of the phenotypic variance, σg is the standard deviation of the genotypic variance, and µ is the phenotypic mean (Falconer and Mackay, 1996).

### Population structure and association analysis

Structure analysis and stratification of the studied population into subpopulations with different genetic structures was done based on SRAP marker data in STRUCTURE software version 2.3.4 (Pritchard *et al*., 2000). This analysis was performed applying an admixture model, a burn-in of 10,000 iterations followed by 100,000 Monte Carlo Markov Chain (MCMC) replicates. The membership of each genotype was run for the range of genetic clusters (K) from K= 2 to K= 10 with five repetitions for each K. Delta k approach by Evanno et al. (2005) was used to determine the optimum number of sub-populations, using STRUCTURE HARVESTER (Earl and VonHoldt, 2012).

Association analysis was run by MLM (Yu *et al*., 2006) to calculate *P*-values for marker–trait associations, using TASSEL version 4.2.1 (Bradbury *et al*., 2007). The phenotypic mean of traits (P-matrix) was applied to identify significant associations under normal and water stress environments, separately. To correct for population structure in MLM model, the Q-matrix derived from structure analysis (at maximum DK), was used as a covariate. Moreover, a kinship matrix (K-matrix) was calculated based on the results of marker genotype data using TASSEL version 4.2.1 (Bradbury *et al*., 2007) and was used. A correction for multiple testing was performed with the FDR (false discovery rate) method, using the QVALUE R package (Storey, 2002).

## Results

### Phenotyping

Mean comparisons of all measured traits for the two moisture environments are given in Table 1. Results showed that moisture environment had significant effect on all of the evaluated traits and the magnitude of mean performance was significantly decreased for all traits under water stress environment (Table 1). Under water stress, DMY1 was approximately reduced by 36, 39 and 37% during 2013, 2014, and 2015, respectively, when compared with normal environment. For DMY2, these reductions were approximately 38, 60, and 56%, in the same three consecutive years relative to normal environment.

**Table 1 -.** Mean performance, phenotypic coefficient of variation (PCV), genotypic coefficient of variation (GCV) and broad-sense heritability (h^2^ _b_) of traits recorded under normal and water stress environments in smooth bromegrass genotypes.

| Traits | Mean ± SD | | Change (%) | PCV (%) | | GCV (%) | | $h^2_b$ (%) | | |
| --- | --- | --- | --- | --- | --- | --- | --- | --- | --- | --- |
|  | Normal | Stress |  | Normal | Stress | Normal | Stress | Combined | Normal | Stress |
| DMY1-Y1 (g/plant) | 155.66 ± 41.77 | 100.24 ± 33.96 | 35.60** | 26.00 | 33.32 | 22.59 | 29.27 | 73.72 | 75.53 | 77.19 |
| DMY2-Y1 (g/plant) | 30.48 ± 6.36 | 18.81 ± 5.42 | 38.29** | 31.05 | 28.80 | 27.35 | 24.18 | 75.93 | 77.61 | 70.50 |
| DMY1-Y2 (g/plant) | 264.59 ± 68.05 | 161.33 ± 50.46 | 39.03** | 24.81 | 31.16 | 22.11 | 27.48 | 70.20 | 79.38 | 77.81 |
| DMY2-Y2 (g/plant) | 61.76 ± 17.19 | 24.81 ± 7.86 | 59.83** | 27.62 | 31.57 | 22.50 | 26.80 | 60.53 | 66.39 | 72.05 |
| DMY1-Y3 (g/plant) | 155.56 ± 42.81 | 98.40 ± 28.16 | 36.74** | 27.32 | 28.34 | 22.98 | 24.42 | 72.69 | 70.77 | 74.27 |
| DMY2-Y3 (g/plant) | 36.92 ± 9.31 | 16.29 ± 4.55 | 55.88** | 25.44 | 27.96 | 19.87 | 22.26 | 59.58 | 60.99 | 63.34 |
| RY (g/plant) | 39.77 ± 11.10 | 25.90 ± 16.75 | 34.87** | 37.29 | 63.43 | 27.14 | 55.10 | 50.93 | 52.97 | 75.47 |
| DRAD (0-9) | 5.45 ± 1.08 | 3.19 ± 1.62 | 41.47** | 19.87 | 50.43 | 14.71 | 44.66 | 54.37 | 54.79 | 78.46 |
| PER (g/plant) | 69.27 ± 49.80 | 49.66 ± 35.14 | 28.31** | 87.24 | 68.46 | 68.15 | 52.49 | 76.33 | 61.03 | 58.78 |
| SDI-Y1 | 8.60 ± 0.34 | — | — | 3.82 | — | 2.86 | — | — | 56.25 | — |
| SDI-Y2 | 7.61 ± 0.56 | — | — | 7.30 | — | 5.86 | — | — | 64.57 | — |
| SDI-Y3 | 7.44 ± 0.74 | — | — | 9.40 | — | 7.96 | — | — | 71.76 | — |

\*\* significant at the 0.01 probability level; ns: not significant

Phenotypic coefficient of variation (PCV) was from 3.82% for SDI-Y1 to 87.24% for PER under normal environment and from 27.96% for DMY2-Y3 to 68.46% for PER under water stress environment. The range of genetic coefficient of variation (GCV) was from 2.86% for SDI-Y1 to 68.15% for PER under normal environment and from 22.26% for DMY2-Y3 to 55.10% for RY under water stress environment. Except for DMY2-Y1, DMY1-Y3, and PER for PCV, and DMY2-Y1 and PER for GCV, the values of genetic variation under water stress were higher than the ones for normal environment (Table 1).

Broad-sense heritability estimates for combined data ranged from 50.93% for RY to 76.33% for PER. Moreover, the heritability estimates were calculated for each moisture environment, separately and are given in Table 1. Under both moisture environments, moderate to high values of heritability were estimated for all traits. According to the results, heritability estimates ranged from 52.97% for RY to 79.38% for DMY1-Y2 under normal environment and from 58.78% for PER to 78.46% for DRAD under water stress environment (Table 1).

### Population structure and association analysis

The maximum likelihood and D*K* were used to calculate the number of subpopulations (*K*). The optimum number of sub-populations (K) was determined based on 30 SRAP primer combinations using the largest value of Delta K in the STRUCTURE 2.3.4 software. The maximum value of D*K* obtained at *K*= 5, suggesting that there is five subpopulations in the smooth bromegrass panel (Supplementary Table S2; Figs. 1 and 2).

**Fig. 1 -.**
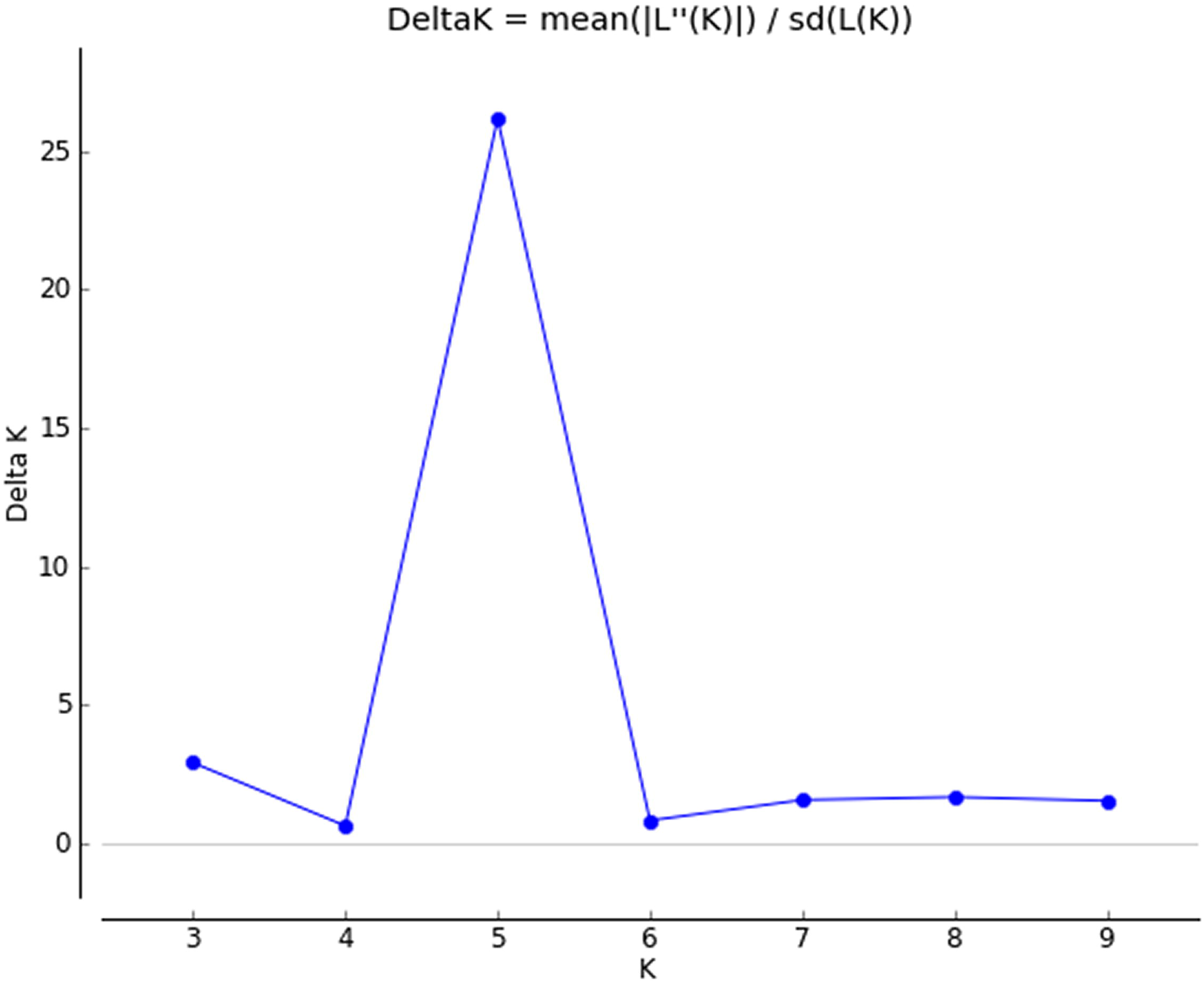
Population structure analysis in a diversity panel of smooth bromegrass genotypes (Δk was used to determine the optimum k value for population structure using the Bayesian clustering method).

**Fig. 2 -.**
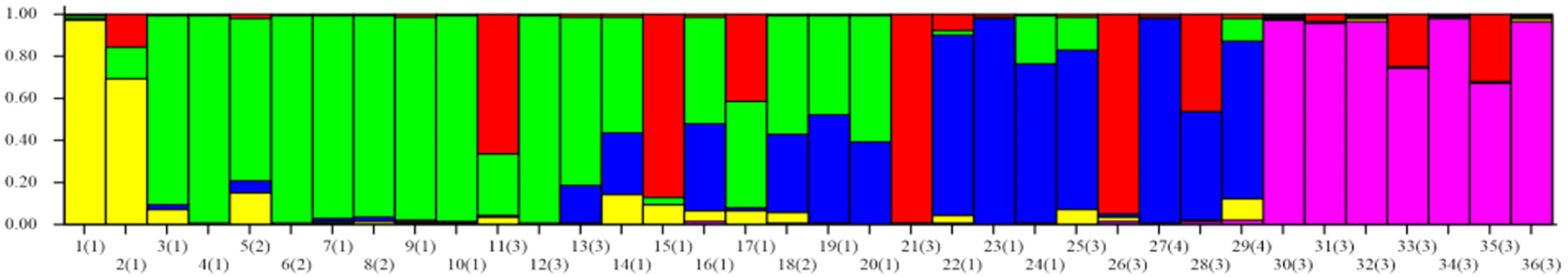
Genetic relatedness of smooth bromegrass genotypes analyzed by STRUCTURE program. Numbers on the *y*-axis indicate the membership coefficient. The color of the bar indicates the five sub-groups identified through the STRUCTURE program. Genotypes with the same color belong to the same group.

Association analysis between SRAP markers and the phenotypic mean of traits was separately conducted for normal and water stress environments, based on MLM model. In this model, kinship or relatedness matrix was considered as a factor. Under normal environment (*P* values <0.01 and a cut-off value of 0.05 for the FDR) 267 and 72 SRAP markers showed significant associations with means of the studied traits, at 0.05 and 0.01 probability levels, respectively (Table 2). The percentage of phenotypic variation (coefficient of determination, *R*^2^) of an individual trait explained ranged from 1.63% to 11.38% (Table 2). Under water stress 185 and 48 markers had significant association with the studied traits, at the 0.05 and 0.01 probability levels, respectively. The percentage of phenotypic variation (*R*^2^) of a trait explained varied from 1.03% to 16.03% (Table 2). It should be stated that, from SRAP markers which were associated with studied traits under normal environment, 72 and 22 markers were significantly associated with SDI-Y1, SDI-Y2, and SDI-Y3 (which were only calculated under normal environment), at the 0.05 and 0.01 probability levels, respectively (Table 2).

**Table 2 -.**
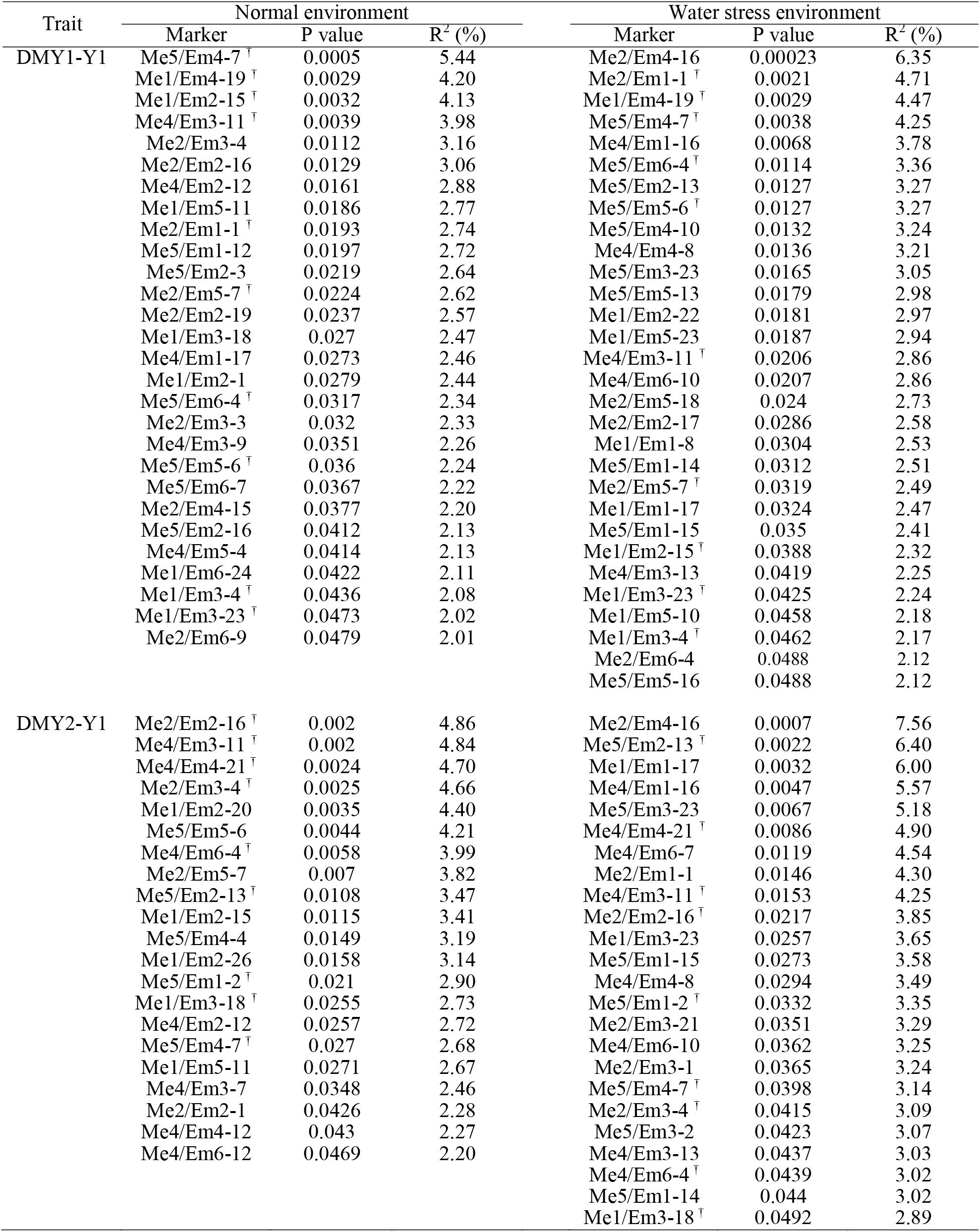
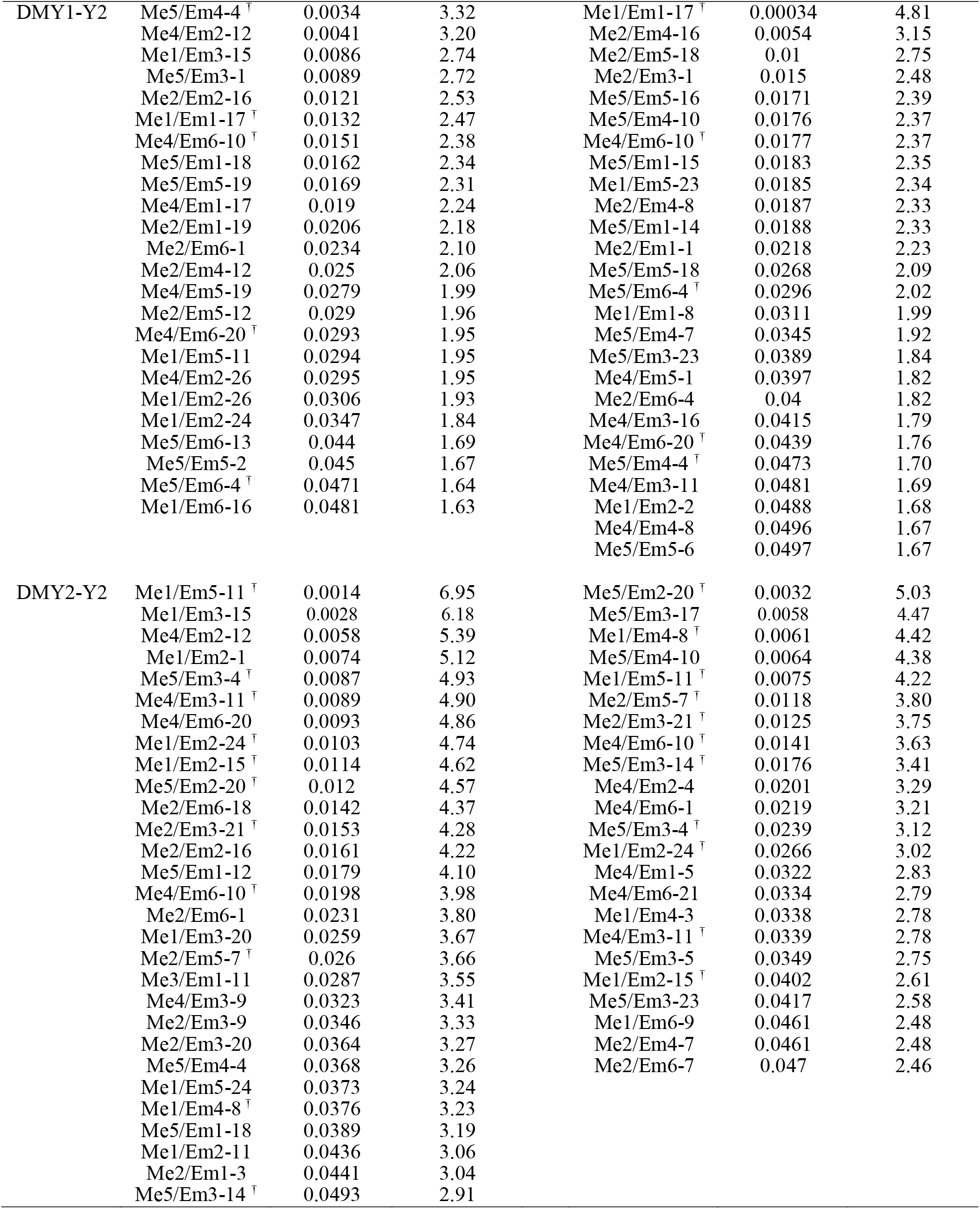
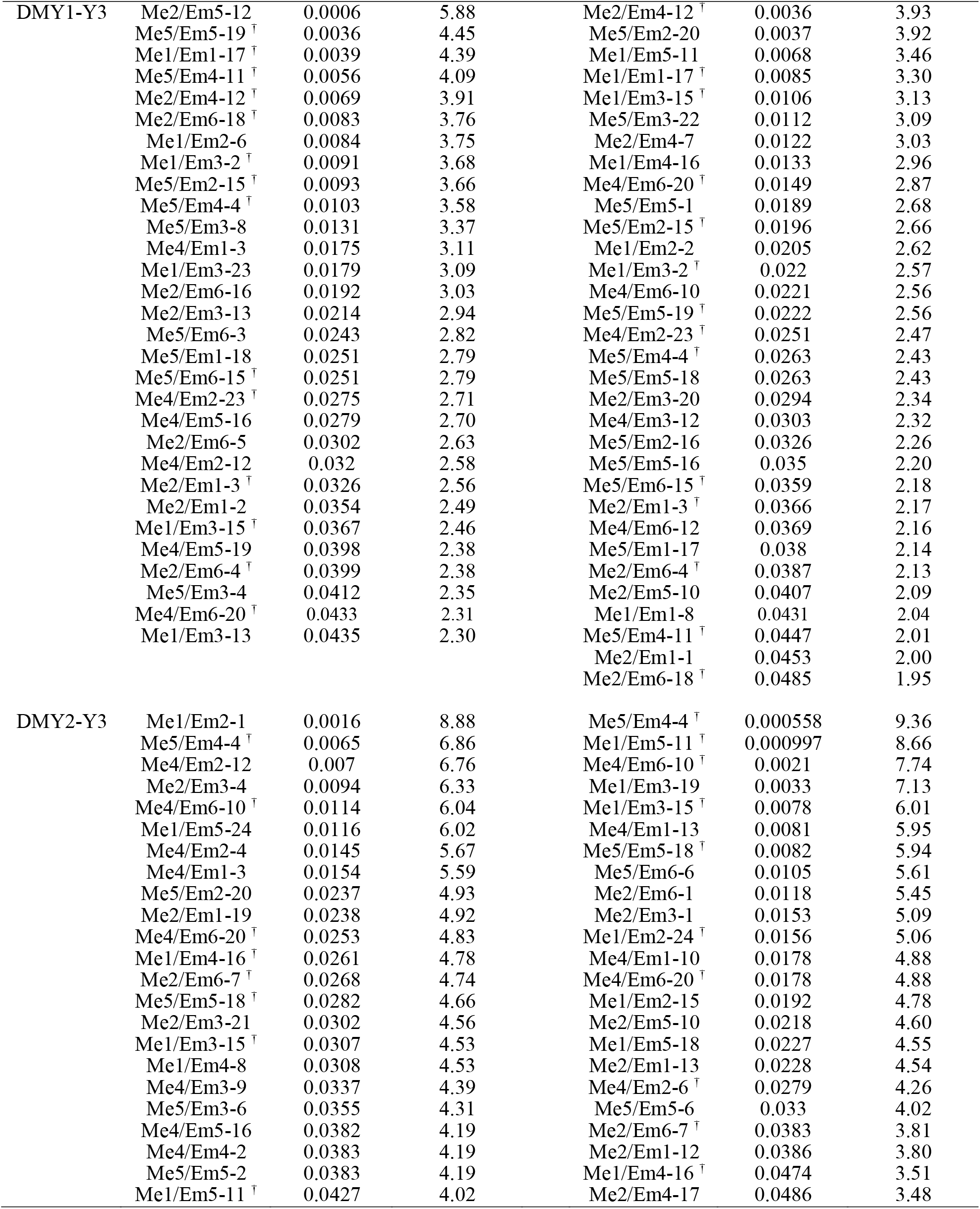
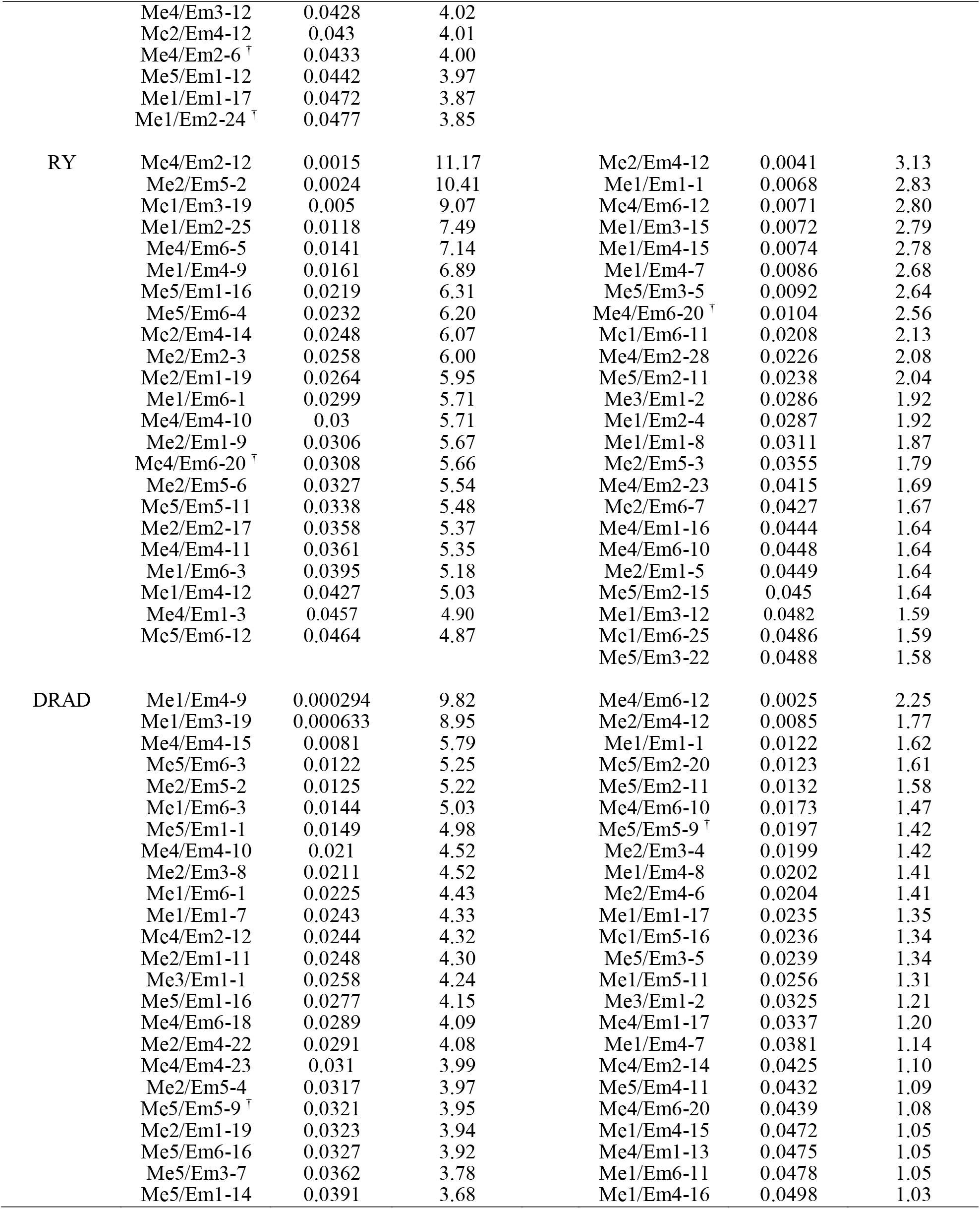
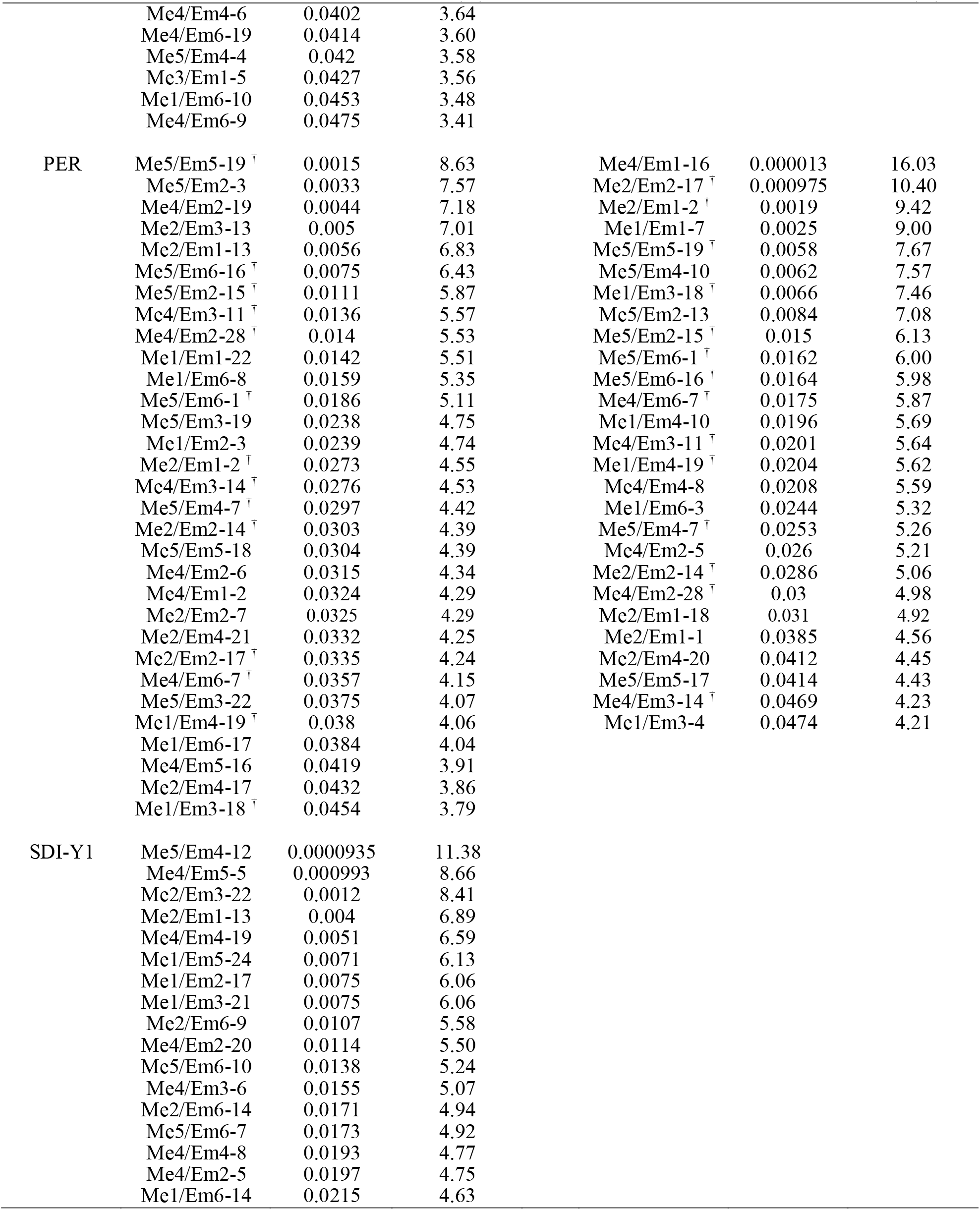
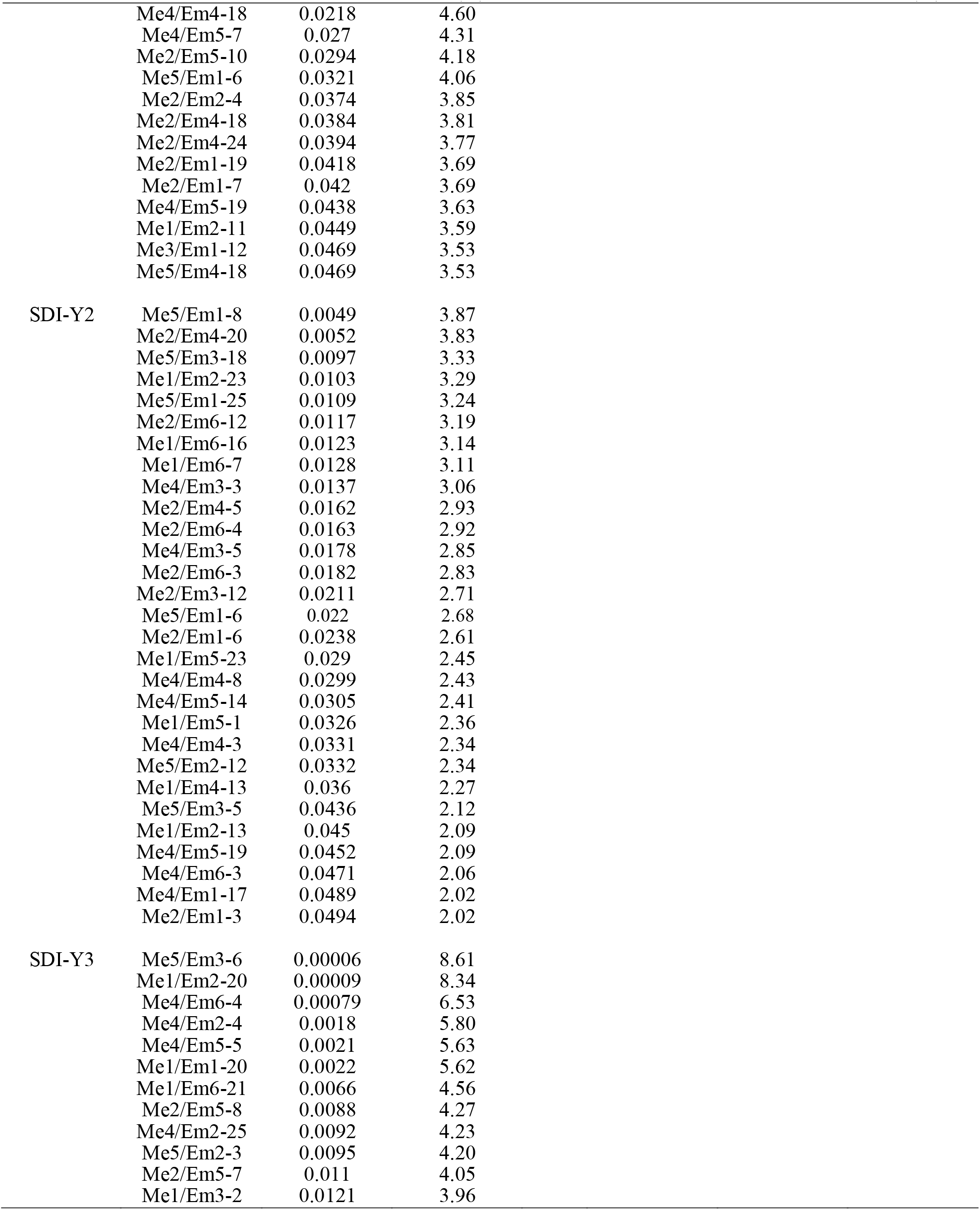
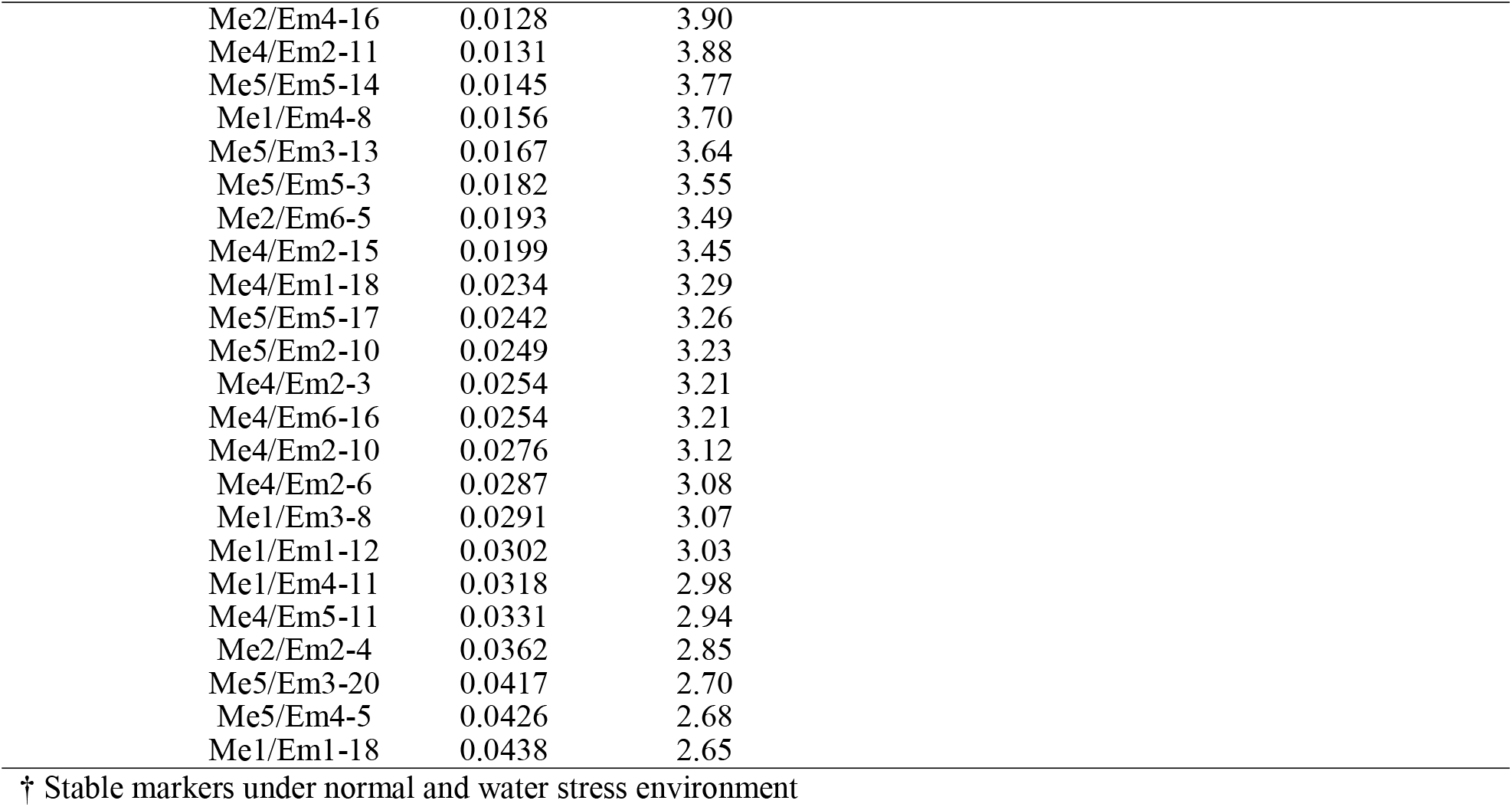
Association of SRAP markers with dry matter yield, persistence, summer dormancy and recovery traits of smooth bromegrass under normal and water stress environments based on mixed linear model (MLM).

Association analysis based on MLM model showed markers which were associated with more than one trait at the same time. Under normal environment 72 markers and under water stress 61 markers showed significant associations with more than one trait, simultaneously. For example, under normal environment, marker Me4/Em2-12 showed significant associations with DMY1-Y1, DMY2-Y1, DMY1-Y2, DMY2-Y2, DMY1-Y3, DMY2-Y3, RY, and DRAD and marker Me4/Em6-20 had significant associations with DMY1-Y2, DMY2-Y2, DMY1-Y3, DMY2-Y3, and RY concurrently. In addition, under water stress situation, marker Me4/Em6-10 showed significant associations with DMY1-Y1, DMY2-Y1, DMY1-Y2, DMY2-Y2, DMY1-Y3, DMY2-Y3, RY, and DRAD and marker Me4/Em6-20 had significant associations with DMY1-Y2, DMY1-Y3, DMY2-Y3, RY, and DRAD (Table 2).

Association analysis was performed in each moisture environment separately to assess stable associations. In total, 75 trait associated markers showed sufficiently stable expression across moisture environments. For instance, markers Me5/Em4-4, Me1/Em1-17, Me4/Em6-10, Me4/Em6-20, and Me5/Em6-4 showed significant and stable associations with DMY1-Y2 in both moisture environments. Similarly, markers Me5/Em4-7, Me1/Em4-19, Me1/Em2-15, Me4/Em3-11, Me2/Em1-1, Me2/Em5-7, Me5/Em6-4, Me5/Em5-6, Me1/Em3-4, and Me1/Em3-23 showed significant and stable associations with DMY1-Y1 (Table 2). Moreover, under both moisture environments, Marker Me1/Em4-19 had significant associations with DMY1-Y1 and PER. In the same way, marker Me5/Em4-7 was associated with DMY1-Y1, DMY2-Y1, and PER under normal environment and with DMY1-Y1, DMY2-Y1, DMY1-Y2, and PER under water stress environment (Table 2).

For assessing the stability of marker-trait associations (MTAs) between years, analysis was conducted on the traits of each experimental year. Results revealed 30 and 28 stable MTAs between years, under normal and water stress environments, respectively (Table 2). For example, under normal environment marker Me1/Em5-11 showed significant associations with DMY1-Y1, DMY2-Y1, DMY1-Y2, DMY2-Y2, and DMY2-Y3. Similarly, marker Me4/Em5-19 had significant and stable associations with DMY1-Y2, DMY1-Y3, SDI-Y1, and SDI-Y2, under normal environment. On the other hand, two markers of Me2/Em1-1 and Me1/Em1-17 showed significant and stable associations with DMY1-Y1, DMY2-Y1, DMY1-Y2, and DMY1-Y3, under water stress environment.

## Discussion

The quantitative inheritance of drought tolerance and interaction between gene expression and environment has challenged the understanding of genetic basis of drought tolerance-related traits in plants (Sun *et al*., 2015). In the present study significant genetic variations were observed among genotypes in terms of all measured traits demonstrating the difference in genes controlling these traits. The non-static performance of genotypes in the two moisture environments emphasizes the importance of marker-trait association analysis in the two moisture environments, separately.

All traits were significantly affected by water stress more likely due to decreased water potential of the soil and decline in net assimilation and photosynthesis of leaves (Flexas *et al*., 2004; Merewitz *et al*., 2010). Saeidnia et al. (2017b) in smooth bromegrass (*Bromus inermis*) and Majidi et al. (2016) in orchardgrass (*Dactylis glomerata*) reported similar results. Wide genetic variation observed for all studied traits revealed potential for genetic gain from selection in this germplasm. Moreover, higher estimates of PCV and GCV for most of the evaluated traits under water stress compared with normal environment indicates that water stress have increased genetic variation and therefore, selection under this condition would be more effective. The findings in this context are contradictory. For example, some researchers believe that genetic gain through selection is higher under normal environment than water limited conditions (Blum, 2011; Majidi *et al*., 2016). While, others have reported higher genetic advance through selection under water stress (Abtahi *et al*., 2018a; Saeidnia *et al*., 2017a). Moderate to high heritability estimates observed for all of the traits emphasizes that detecting of marker–trait associations is possible for these traits (Hung *et al*., 2012).

Population structure analysis separated the smooth bromegrass genotypes into five groups with different genetic structures. Furthermore, association analysis of evaluated traits under normal and water stress conditions revealed that MTAs were mostly different at the two environments. Moreover, results showed that a greater number of genes were involved in controlling traits under normal environment than water stress. The percentage of variation which is explained by each identified association was low, that may be due to the role of many minor genes controlling the trait, outcrossing nature of smooth bromegrass, markers exhibiting minor quantitative effect, rare alleles, and complex allelic interactions (Yang *et al*., 2010; Debibakas *et al*., 2014). Similar results were reported by Lou et al. (2015) and Sun et al. (2015 in tall fescue.

Based on the results, some markers had simultaneously significant associations with more than one trait and may be effectively used for the improvement of several traits, concurrently (Sun *et al*., 2015; Abtahi *et al*., 2018b). Association between multiple traits could be attributed to the co-expression mediated by expression of quantitative trait loci or e-QTLs (House *et al*., 2014). For instance, marker Me2/Em1-19 showed simultaneously significant associations with DMY1-Y2, DMY2-Y3, RY, DRAD, and SDI-Y1 under normal environment. Similarly, marker Me4/Em3-11 concurrently showed significant associations with DMY1-Y1, DMY2-Y1, DMY1-Y2, DMY2-Y2, and PER, under water stress environment. The simultaneous associations of markers with multiple traits have been attributed to the pleiotropic effects or to several tightly linked genes affecting the traits (Lehner, 2011; Sun *et al*., 2015).

Improving the persistence, recovery, and survivability of perennial forage grasses is one of the main objectives of grass breeders in areas with prolong periods of drought (Annicchiarico *et al*., 2011; Pecetti *et al*., 2011). In this study, markers associated with these traits were identified under normal and water stress environments. For instance, markers Me5/Em5-19, Me5/Em6-16, Me5/Em2-15, Me4/Em3-11, Me4/Em2-28, Me5/Em6-1, Me2/Em1-2, Me4/Em3-14, Me5/Em4-7, Me2/Em2-14, Me2/Em2-17, Me4/Em6-7, Me1/Em4-19, and Me1/Em3-18 were associated with superior persistence under both moisture environments. Similarly, marker Me4/Em6-20 was associated with RY, and marker Me5/Em5-9 was associated with DRAD, under normal and water stress environments. If the effectiveness of these regions in the genetic control of these traits is confirmed, these markers could be potentially used for the improvement of recovery, persistence and therefore drought tolerance of smooth bromegrass. Moreover, an important trait associated with survivability, persistence and recovery after drought is summer dormancy; which improves autumn recovery and therefore results in a better persistence of perennial grasses. In the present study 30, 29, and 35 markers were associated with SDI-Y1, SDI-Y2, and SDI-Y3, respectively. From these markers, 24 were also associated with other traits and four markers showed stable association and were associated with SDI at two or three years of study.

In the present study most of the MTAs were different for the two moisture environments and also for the three experimental years, indicating the considerable role of environmental effects in these associations (Bocianowski and SeidlerŁozykowska, 2012). These findings may suggest that different genes contribute to the same trait in different environments and years (Rumbaugh *et al*., 1984) or the same genes may change the expression level between different environments (House *et al*., 2014). In the present study, 75 markers showed stable association with different traits under both moisture environments. Moreover, 30 and 28 markers showed stable MTAs between three years of study, under normal and water stress environments, respectively. In general, associated markers which were detected in two or more different environments or experimental years are more reliable than those present in only one environment (Diapari *et al*., 2015).

In conclusion, the advantage of association analysis technique as a powerful tool to identify and detect genes and markers linked to complex traits of agricultural and economic importance was demonstrated. Satisfactory levels of polymorphism and genetic diversity was observed for the studied traits in the polycrossed population. Five subpopulations were identified in smooth bromegrass panel; and 339 and 233 significant MTAs were detected under normal and water stress environments, respectively. Some SRAP markers were associated with the recovery and persistence of this species. Therefore, it was demonstrated that SRAP markers can be used in the future breeding program to improve recovery after prolonged drought and persistence, and hence enhance drought tolerance of smooth bromegrass. Environmental specificity of MTAs indicated that genotype × environment interactions affect association analysis; nevertheless, 75 MTAs showed significantly stable expression across normal and water stress conditions. Also, 30 and 28 MTAs showed significantly stable expression across years of study, under normal and water stress conditions, respectively. The molecular markers identified in the present study are suggested as useful genomic resources in the future breeding programs of smooth bromegrass.

## Supplementary data

Supplementary data are available at *JXB* online.

*Table S1*. Information on parental plants of genetic materials used in this study.

*Table S2*. Calculated statistics to detect optimum number of subgroups (*K*) in structure analysis of smooth brome genotypes (D*K* method; Evanno *et al*. 2005), using the program STRUCTURE.

## Acknowledgements

We would like to thank *Iran National Science Foundation (INSF)*[grant number 98025737]and Isfahan University of Technology for the award of a Postdoctoral Research Fellowship to the first author.

## Author contributions

FS, MMM and AM conceived and designed the experiments; FS performed the experiments, analyzed the data and wrote the manuscript with the supervision of MMM and AM; all authors discussed the results and reviewed the manuscript.

## Data availability statements

The data supporting the findings of this study are available from the corresponding author, (F. Saeidnia), upon request.

### Abbreviations

DMY1: dry matter yield of cut 1
DMY2: dry matter yield of cut 2
DRAD: degree of recovery after drought
GCV: genotypic coefficient of variation
GLM: general linear model
GWAS: genome wide association study
LD: linkage disequilibrium
MAS: marker assisted selection
MLM: mixed linear model
MTA: marker trait association
PCV: phenotypic coefficient of variation
PER: persistence
QTL: quantitative trait loci
RY: recovery yield
SDI: summer dormancy index
SRAP: sequence related amplified polymorphism.

## Notes

### Competing Interest Statement

The authors have declared no competing interest.

